# Spatial Adaptation of Primate Retinal Ganglion Cells between Artificial and Natural Stimuli

**DOI:** 10.1101/2025.04.09.647910

**Authors:** Michaela Vystrčilová, Shashwat Sridhar, Max F. Burg, Mohammad H. Khani, Dimokratis Karamanlis, Helene M. Schreyer, Varsha Ramakrishna, Steffen Krüppel, Sören J. Zapp, Matthias Mietsch, Tim Gollisch, Alexander S. Ecker

## Abstract

The retina encodes a broad range of stimuli, adapting its computations to features like brightness, contrast, or motion. However, it is unclear to what extent it also adapts to spatial frequency content – as theories of efficient coding would predict – for instance, when switching between natural scenes and white noise. To address this, we analyzed neural activity of marmoset retinal ganglion cells (RGCs) in response to white noise and naturalistic movie stimuli. We trained linear-nonlinear models on both stimuli, evaluated their performance and compared their receptive fields (RFs) across the stimulus domains. We found that the models with spatial filters trained on either one of the stimulus ensembles were not able to predict the neural activity on the other as accurately as the models trained on the target stimulus. This suggests that spatial processing adapts to stimulus statistics. Different RGC types exhibited distinct changes: the midget OFF cells’ RFs became enlarged under natural movie statistics, resulting in a lower cutoff frequency. Parasol cells did not change their RF size significantly. Large OFF cells’ RFs decreased in size. All cell types exhibited stronger surrounds under natural movies, resembling the whitening filters predicted by efficient coding. However, quantifying the effect of the filter adaptation on the stimulus power spectrum showed a significant contribution towards whitening only in ON parasol cells. The whitening effect emerged regardless of the training stimulus. These results suggest that while RGCs adapt to the spatial frequency content of the input, efficient coding can only partially account for this adaptation.

**Significance statement:** Natural scenes differ from artificial stimuli like white noise, in spatial frequency structure. How the retina adapts to these differences remains unclear. To explore this, we studied responses of four primate retinal ganglion cell types to artificial and natural stimuli. Our results show that some cell types, like midget cells, enlarge their receptive fields and surrounds under natural stimuli, while others, like parasol cells, only enhance surrounds. These changes align qualitatively with the efficient coding theory, which posits redundancy reduction. However, in three cell types, the enhanced surrounds did not significantly whiten responses to natural stimuli, contrary to efficient coding predictions. These findings challenge how fully efficient coding explains retinal adaptation, suggesting that other principles underlie the processing of visual inputs.

## Introduction

The retina is the direct interface between the visual environment and an animal’s brain. It evolved to encode stimuli with diverse statistical properties by dynamically adjusting to specific attributes of visual inputs. Adaptation to the mean luminance (Dowling, 1960; Dowling, 1963) and contrast levels (Smirnakis et al., 1997; Kim and Rieke, 2001; Baccus and Meister, 2002) is well documented, while higher-order statistics (skew, kurtosis) showed only mild influence (Tkačik et al., 2014). Salamander and rabbit RGCs adapt to spatiotemporal correlations in the stimulus, such as temporal correlations in flickering light intensity, anti-correlated spatial patterns in checkerboard stimuli and grating orientations (Smirnakis et al., 1997; Hosoya, Baccus, and Meister, 2005). Adaptation to motion statistics has been shown using moving gratings (Ölveczky, Baccus, and Meister, 2007) and parameterized motion clouds (Ravello et al., 2019).

Given the many reported forms of adaptation, one might expect neurons to also adapt to the spatial frequency content of the stimulus. This is indeed the case in later stages of the visual system. For instance, cat V1 simple cells have significantly different spatial RFs under white noise and natural image stimulation (Sharpee, Miller, and Stryker, 2008). In the cat’s LGN, Lesica et al. (2007) found that under natural movie stimulation the cells’ RFs were spatially larger and had a more pronounced surround, compared to white noise stimulation. In the retina, Nirenberg et al. (2010) looked at the temporal adaptation of RGCs’ RFs between white noise and natural movies, and found different levels of adaptation between ON and OFF cells. However, it is unknown whether the spatial structure of RGCs’ receptive fields adapts to the different statistics of white noise and natural scenes.

Many properties of the visual system have been explained by the efficient coding hypothesis, including the RF structure (Doi et al., 2012; Atick and Redlich, 1992; Gupta et al., 2023), the ratio of ON and OFF cells (Ratliff et al., 2010), and the emergence of cell types and their mosaic arrangement (Karklin and Simoncelli, 2011; Ocko et al., 2018; Jun, Field, and Pearson, 2022; Roy et al., 2021). Efficient coding posits that neuronal processing minimizes redundant information (Barlow, 1961; Linsker, 1988; Atick and Redlich, 1990). Lesica et al., 2007 show that LGN neurons adapt their RF surround strength and center size in response to white noise and natural movies in line with efficient coding principles: a stronger surround whitens the correlated movie stimulus and an increased center size enhances low-pass filtering to remove low-power frequencies in natural movies.

Given the optic nerve’s limited capacity, efficient coding could be particularly applicable to the retina. Stimulus adaptation could facilitate more efficient information transmission through this bottleneck. However, it is currently unknown how the spatial RFs of RGCs adjust to spatial frequency content, whether this adaptation follows the predictions of efficient coding and how this adaptation manifests across cell types. Addressing this gap is crucial for understanding the foundational mechanisms of visual processing and the extent to which the retina contributes to the changes we observe in the LGN.

To answer these questions, we investigate the adaptation in different types of RGCs in the marmoset retina, training linear-nonlinear (LN) models (Marmarelis and Naka, 1972; Korenberg and Hunter, 1986; Sakai, Ken-Ichi, and Korenberg, 1988; Chichilnisky, 2001; Pillow et al., 2008) directly on different stimuli types and comparing the resulting model parameters. We show that models perform best when tested on the same stimulus statistics they were trained on, and performance deteriorates when testing on the other stimulus type, suggesting that RGCs RFs adapt to stimulus statistics.

Furthermore, we find that midget cells exhibit larger center sizes and stronger RF surrounds when responding to naturalistic movies compared to white noise. In contrast, parasol and large OFF cells only display stronger surrounds. Although many of these changes are qualitatively in line with the predictions of efficient coding about stimulus adaptation, a quantitative analysis of filtered stimulus spectra reveals that the surrounds of OFF midget, OFF parasol and large OFF cells, contribute only minimally to whitening of the natural movie signal. In ON parasol cells, we do observe a significant whitening effect of RFs estimated from both white noise and natural movies. While the whitening in parasol cells can be interpreted as following the efficient coding theory, together, these findings suggest that additional principles are required to account for retinal adaptation.

## Methods

### Data acquisition and stimuli

We used electrophysiological recordings of the responses of retinal ganglion cells (RGCs) to artificial and naturalistic stimuli, obtained from adult common marmosets (*Callithrix Jacchus*). The acquisition and preprocessing of this data has been described in detail in Sridhar et al. (2024). In brief, the retina was extracted from the enucleated eye and mounted on a micro-electrode array (MEA) while under constant perfusion with an oxygenated solution and while ensuring minimal light contamination of the tissue. A monochromatic oLED screen, focused on the photoreceptor-layer of the mounted tissue, was used to project the stimuli while the activity of RGCs was recorded using the MEA setup. Two types of stimuli were presented: spatiotemporal white noise and a naturalistic movie overlaid with simulated eye movements. The resolution of the stimuli was 600×800 pixels, naturalistic movie was shown at full resolution and the white noise was projected at 150×200 stimulus pixels, each stimulus pixel corresponding to 4×4 screen pixels and each screen pixel to 7.5×7.5 µmeters. Spiking responses of individual RGCs were isolated using a semi-automated spike-sorting pipeline and binned at the resolution of the stimulus refresh rate of 85 Hz for further modeling.

The stimuli were displayed in repeats – trials, containing non-repeating frames which we used for training and validation and repeating sequences of frames which we used for testing. We used responses of two retinas, data from one retina being one dataset. Both of the datasets contained 10 trials of spatiotemporal white noise stimulus, each consisting of 150 s of non-repeating frames and 30 s of repeating frames and 20 trials of the naturalistic movie, each consisting of 300 s of non-repeating frames and 60 s of repeating frames.

### Cell selection and classification

The recorded cells were screened on the basis of the reliability of their responses to both stimuli and then assigned cluster labels based on their receptive field and response properties, as described in detail in Sridhar et al. (2024). The cells’ reliability was measured using the fraction of explainable variance (Cadena et al., 2019) and the symmetrized coefficient of determination (Karamanlis and Gollisch, 2021). Four cell clusters were isolated based on receptive field and response properties of the cells – OFF midget, OFF parasol, ON parasol and large OFF. While the first three correspond to known cell types in the primate retina, the large OFF cells were composed of a mixture of cell clusters with no known cell type correlates in primates. This left us with 172 reliable, classified cells across two datasets – *Dataset 1* (98 cells) and *Dataset 2* (74 cells).

### Linear-nonlinear (LN) models

We trained two types of LN models, rank-one and rank-two. Rank-one models were consisted of a separable spatial and temporal filter, as an estimate for a cell’s RF, with each spatial filter cropped to 15×15 pixels around the center of the spatial RF. The rank-two LN models consisted of two such separated spatial and temporal filter pairs with the same center and crop sizes as rank-one models. The sum of the outer products of both of the filter pairs was the estimate for a cell’s RF.

The pixel with the largest temporal variance in the spatial RF estimate obtained from spike-triggered averaging on responses to spatiotemporal white noise was the RF center. We calculated the STA as

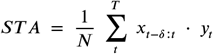

where *x*_*t*_ is the frame presented at time *t, y*_*t*_ is the spike-count recorded at time-point *t* and *N* is the recorded number of spikes and *T* is the total number of frames. The temporal filter length was 30 frames for both rank-one and rank-two LN models, corresponding to a time window of about 350 ms. To fit each LN model, the stimulus frames were also cropped to 15×15 pixels around the RF center of the cell and then linearly convolved with the spatial and temporal filter before passing through a non-linearity. The non-linearity was a softplus function with two learnable parameters, *α* and *β*:

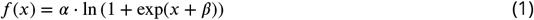

We learned the spatial and temporal filters along with the parameters of the non-linearity endto-end during model training.

### Training

All parameters of the LN model were optimized with stochastic gradient descent using the following training scheme. We shuffled the non-repeating parts of the stimulus trials and assigned 80% of them to the training set and the remaining 20% to the validation set. We randomly initialized the filter values and parameters of the output non-linearity, and updated them by minimizing the Poisson loss,

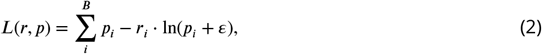

between the predicted *p*_*i*_ and the observed *r*_*i*_ response values for bin *i. ε* was set to 10^−12^ to avoid computing *ln*(0). We trained the models for up to 1000 epochs, with early stopping when the correlation between the recorded and predicted responses in the validation trials had not improved for 30 consecutive epochs. The optimizer we used was Adam (Kingma and Ba, 2017) with an initial learning rate of 0.005. We employed the Pytorch learning rate scheduler ReduceLROnPlateau with a patience of 15 on the validation correlation with minimal learning rate of 0.000001.

### Out of domain adaptation

To adapt the temporal filter and parameters of the non-linearity obtained from one stimulus to the other, we froze the spatial filter (or filters in the case of the rank-two models) and started training using the same procedure as described above, changing only the parameters of the temporal filter(s) and the non-linearity. The initial learning rate was 0.001, all the other scheduler settings were kept the same as when training all parameters of the model. For this analysis, we used the same number of natural movie and white noise frames for both training the complete model on one stimulus and adapting the temporal filter and non-linearity on the other.

### Evaluation

We reported the final performances of the models on the test dataset by averaging the calculated the correlation coefficients for all cells. For a given RGC *c* the correlation was calculated as in Equation (3) between the predictions *p*_*c*_ and the trial averaged firing rates ⟨*r*⟩_*c*_ on the held-out test sequence with length *T*.

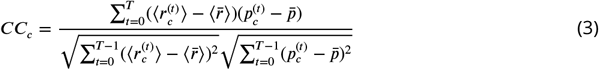

where 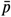 is the within-trial average prediction and 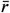 the within-trial averaged response.

### Receptive field size and surround amplitude estimation

For each cell, we obtained the center size and surround amplitude of its rank-one LN-model-estimated RF by fitting a Difference of Gaussians (DoG) to the spatial filter. The DoG was calculated as

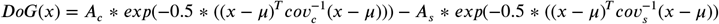

where *A*_*c*_(*A*_*s*_) is the center (surround) amplitude, *μ* the position vector shared by both center and surround and 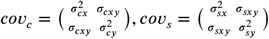 are the covariance matrices of the center (surround) Gaussians in pixel space. The DoG was fitted by minimizing the means squared error between the DoG estimate and the spatial filter. We applied a positivity constraint on the diagonal elements of the covariance matrices of both Gaussians and enforced that both have the same amplitude sign. The initial parameters were *A*_*c*_ = 0.5, *σ*_*cx*_ = 1.2, *σ*_*cy*_ = 1.2, *σ*_*cxy*_ = 10^−3^, *A*_*s*_ = 0.5, *σ*_*sx*_ = 1.5, *σ*_*sy*_ = 1.5, *σ*_*sxy*_ = 10^−3^ and the mean *μ* was set as the center of the crop region. We established the center size *S* as 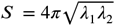. where the *λ*_1_ and *λ*_2_ are the eigenvalues of the center covariance matrix. To obtain the surround strength, the DoG-parameterized spatial filter was normalized to a maximum absolute value of 1. Then, the absolute value of the lowest negative value (if center amplitude was positive) or the highest positive value (if the center amplitude was negative) of the DoG fit was taken to be a measure of the surround strength.

### Mean RF estimation

We normalized the LN model filters such that the maximum of the temporal filter set to 1 (−1) for ON (OFF) cells and averaged across the spatial filters, to calculate the mean RF for each cell type. Before calculating the average, we aligned the centers of all spatial filters within a cell type using the center coordinates from their DoG fits. Empty pixels created due to the center shift were replaced by mean gray.

### Simulated filters

We simulated spatial filters following the principles of efficient coding. First, we simulated filters mimicking those obtained from white noise data i.e. filters with a small center size and no surround. We call these the “white noise” filters. To generate them, we used parameters of the central Gaussian of a DoG fit to the mean midget cell RF, with amplitude normalized to 1 and constructing a diagonal covariance matrix with all diagonal elements equal to the largest diagonal value in the covariance matrix of the DoG fit. This resulted in a circular receptive field of the size of a midget cell with no surround. We followed the same procedure with the mean parasol cell RF to simulate large center sizes and produce a stronger low-pass filtering. We call these the “low-pass” filters.

To simulate the presence of a surround in both, the “white-noise” and the “low-pass” filters, we subtracted a second 2D Gaussian from each. The amplitude and covariance matrix of the second Gaussian were adjusted manually so as to ensure the resulting DoG filter would lead to a flattening of the low-frequency spatial power spectrum for naturalistic movies (see below).

### Power spectra calculation

We calculated the spatial power spectral densities of non-filtered white noise and natural movie stimuli, both at monitor pixel resolution, by performing a 2D Fast Fourier Transform (FFT) on each 800×600 stimulus frame cropped to 600×600 pixels. Radial averaging was applied to the frequency components of each frame, and the results were averaged across frames to obtain the final stimulus power spectra.

To get transfer functions of the learned filters, we padded each filter to match the 600×600 stimulus dimensions and used the same FFT-based procedure, computing individual transfer functions for each LN model’s spatial filter. Cell-type-specific transfer functions were obtained by averaging across filters from the same cell type. Transfer functions for the simulated filters were computed in the same way directly from the simulated filters.

We calculated the spectrum of the natural movie stimulus filtered with both rank-one and ranktwo estimated RFs from both white noise and natural movies. We followed the same procedure in all the settings. We first convolved the natural movie stimulus with each cell’s estimated RF. We then computed the spatial power spectra of the filtered stimulus frames and averaged the spectra of these frames to get cell-specific spectra. To get the spectrum for a given cell type, we averaged over the spectra of cells classified as such. For the simulated filters, we multiplied their transfer functions with the power spectra of each stimulus frame and then averaged over the frames.

## Results

We analyzed the activity of marmoset (*Callithrix jacchus*) RGCs in response to white noise (WN) and naturalistic movie (NM) visual stimuli recorded using micro-electrode arrays (Figure 1). We will also refer to these stimuli simply as “noise” and “movies”. Our datasets contained the activity of 172 RGCs across two marmoset retinas, characterized by Sridhar et al., 2024 which were shown both movie and noise stimuli. Sridhar et al., 2024 classified these cells into four groups, *OFF Midget, OFF Parasol, ON Parasol and Large OFF* which we also used, and excluded cells that responded unreliably (for further details, see Methods). The Large OFF group was an inhomogeneous class of neurons that contained multiple cell clusters with spatially large RFs and consistent response characteristics but without known cell type correlates in primates.

**Fig. 1.**
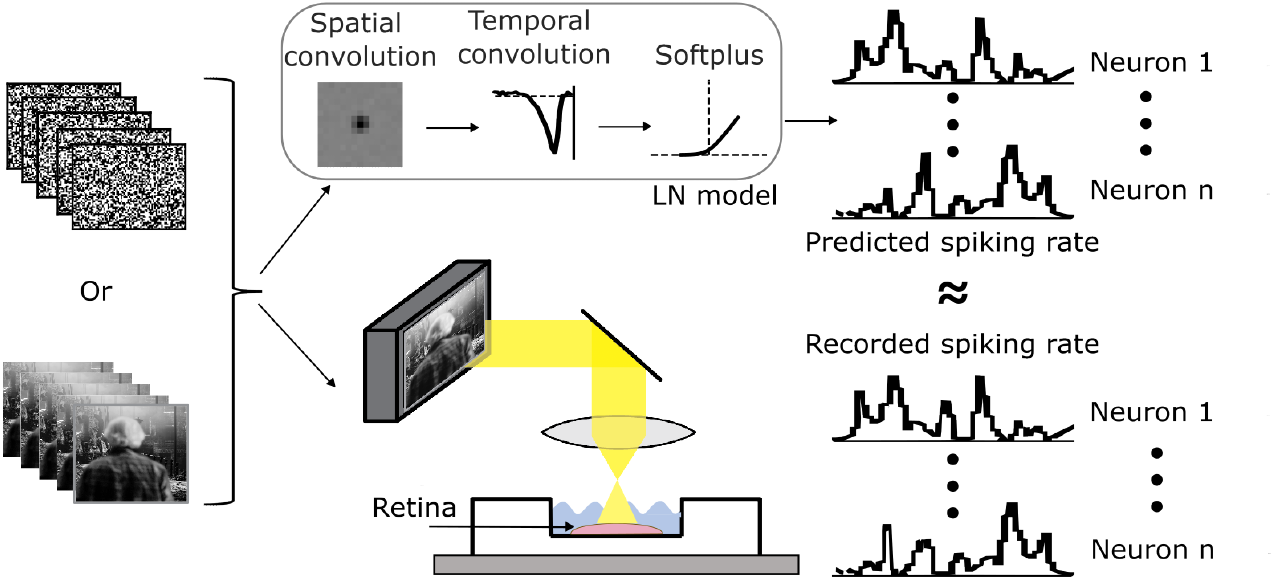
We presented white noise (WN) or naturalistic movie (NM) stimuli to a retina while recording ganglion cell responses using a micro-electrode array. We train end-to-end LN models to predict the average firing rate of all neurons given the stimulus in a time window preceding the prediction.

We split the stimuli into non-repeating, unique video sequences for training models (“training set”) and into a smaller number of repeated video segments for evaluating the final model performance (“test set”). A sequence of a non-repeating segment followed by a repeated segment comprised a “trial”. In our data we showed 10 to 20 stimulus trials to the retina. The length of a non-repeating segment was 150–300 seconds and of the repeating segment 30–60 seconds depending on the stimulus type.

We trained LN models to predict the neural responses in these datasets (Figure 1). To evaluate their performance, we computed the correlation coefficient between the predicted spike counts and trial-averaged neuronal response, which we measured in 1/85 second time bins (corresponding to the frame rate of 85 Hz) on the test segments (Equation (3)). We held out the averaged test segments during training and trained the models only on non-repeating training set data.

### Training stimulus-specific models improves predictive performance

LN models are a classical way of modeling RGC responses to white noise. We start by establishing their predictive performance on both white noise and natural movie stimuli to verify that they are reliable models of the studied RGCs. The LN models consist of a linear spatiotemporal filter capturing the neuron’s linear RF, followed by a non-linearity. For simplicity, we assumed the filter to be space-time separated. To obtain the predicted spiking responses, we took the inner product between the stimulus and the linear filter and applied a parametrized softplus non-linearity (Figure 1).

We obtained the filters and non-linearity parameters of the LN models by fitting them directly on the training set of the data using maximum likelihood estimation with stochastic gradient descent. We fit models independently for each stimulus type – that is two models per cell, one model for white noise and one for natural movies. First, we evaluated the models *in-domain* (ID), i. e. the models trained on white noise data were evaluated on the white noise test data split and models trained on natural movies were evaluated on the natural movies test data split (Figure 2A). The Pearson correlations coefficient between the recorded and model-predicted RGC activity, mean and standard deviation (SD) across cells of a given cell type, were between 0.36 (±0.1 SD) for large OFF cells and 0.73 (±0.07 SD) for OFF midget cells (OFF parasol: 0.7 ± 0.1 SD; ON parasol: 0.64 ± 0.05 SD). On natural movies, the models performed better for all four cell types, with the mean correlation between 0.52 (±0.07 SD) for large OFF cells and 0.86 (± 0.02 SD) for ON parasol cells (OFF midget: 0.76 ± 0.06 SD; OFF parasol: 0.77 ± 0.09 SD).

**Fig. 2.**
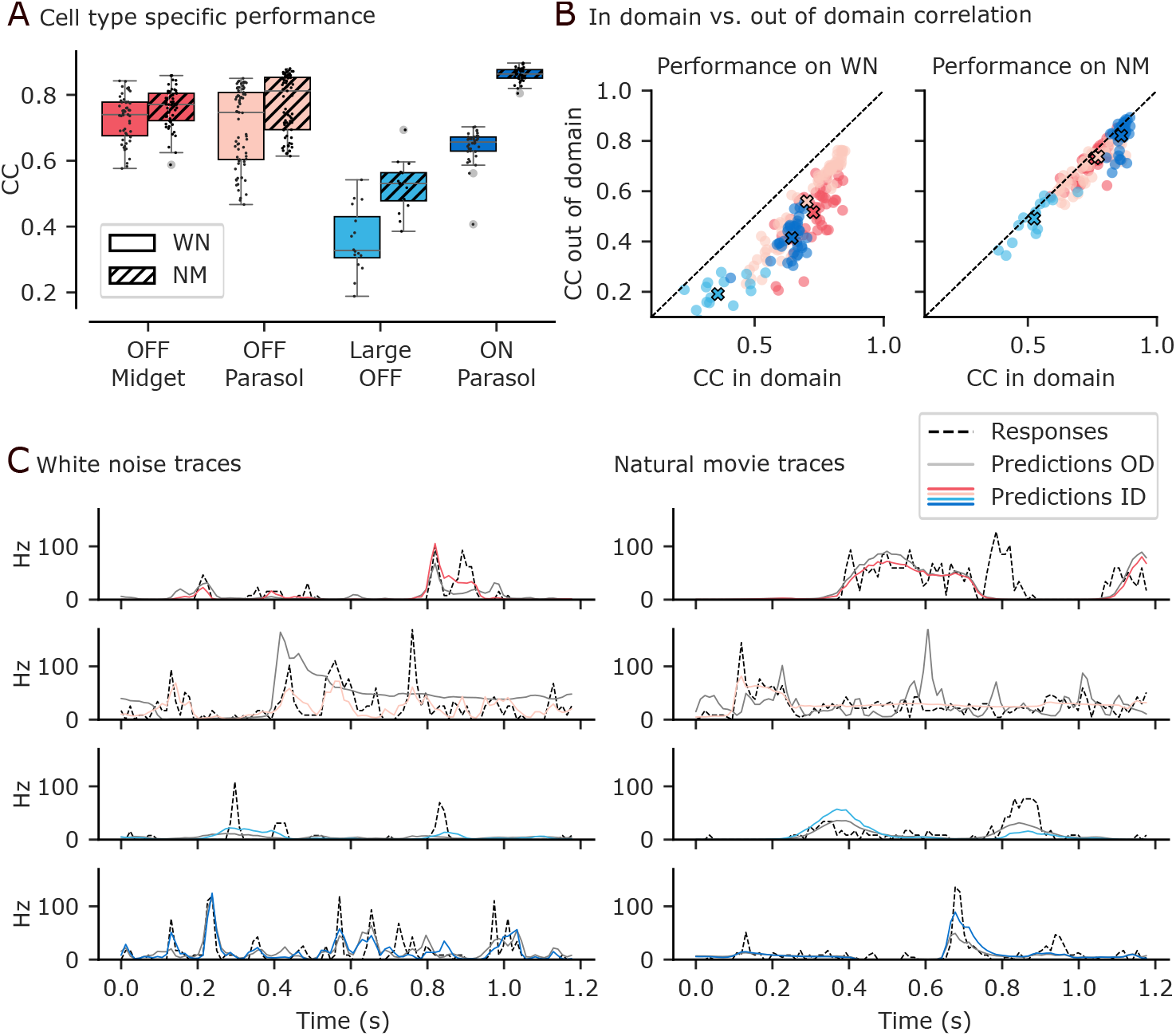
**A**. End-to-end trained LN model correlation coefficient (CC) for the different cell types on white noise (solid) and natural movies (dashed) respectively. **B**. In domain (ID) vs. out of domain (OOD) correlation comparison. Left plot: x axis are models trained and evaluated on WN, y axis are models trained on NM and adapted and evaluated on WN. Right plot: x axis are models trained and evaluated on NM, y axis are models trained on WN and adapted and evaluated on NM. **C**. Comparison of example cells’ prediction for each cell type between recorded signal (black, dashed) in domain predictions (coloured), out of domain predictions (gray).

Next, we asked whether models learned on one stimulus would generalize to the other stimulus – that is, we evaluated their performance *out-of-domain* (OOD). Specifically, we were interested in whether the spatial part of the RF is stimulus-specific or generalizes out-of-domain, i. e. to the other stimulus. Hence, we took a model trained on one stimulus type (for example white noise) and froze its spatial filter weights. We then continued training this model on the other stimulus (natural movies in this example), adjusting only the parameters of the temporal filter and the nonlinearity. Finally, we evaluated the model on the test set of the other stimulus (natural movies). This approach allowed us to determine if the spatial filter generalized across stimuli. If the OOD model’s performance would match the ID performance, it would indicate that the spatial filter is a fixed property of the neuron. If not, it would suggest that the spatial RF adapts, as only the spatial filter was held constant during evaluation.

We found that all models performed better in an in-domain evaluation setting compared to outof-domain evaluation (Figure 2B). The margin in correlation of the ID over the OOD model on white noise was between 0.23 for ON parasol cells and 0.14 for OFF parasol cells (OFF midget: 0.21; large OFF: 0.17). On natural movie, this difference was 0.03 for OFF midget, OFF parasol and large OFF cells and 0.04 for ON parasol cells. This means that spatial RFs of our LN models did not generalize across stimuli and it suggests spatial changes in retinal computation across these two stimuli. The traces of example cells for the different cell types also show that the ID models were able to better follow the recorded firing rates compared to OOD models (Figure 2D).

The white noise models generalized better to natural stimuli than the other way around. Even though the differences between ID and OOD models evaluated on natural movies were small, this does not imply small changes in the RFs between the two stimuli. The natural movie stimuli have most of their energy in the low frequency range, which means that a major fraction of the correlation was explained by a few low frequency components, whereas – as we show below – the RF differences were larger in higher frequencies.

### Retinal ganglion cells adapt their spatial RFs to stimulus statistics

The lack of generalization across the white noise and natural movie stimulus suggests that some features of the RF depend on the type of stimulus. We singled out the generalization of the spatial part of the RF and investigated spatial adaptation of the size and the surround strength of the RFs. First, we investigated the average RF for each cell type, by aligning all RFs to the same location and averaging across neurons of the same type (Figure 3A). We found qualitative effects to differ across cell types. For midget cells, the center RF size increased under natural movie stimulation compared to white noise. For parasol ON and OFF cells, the center size stayed roughly unaffected and for large OFF cells, it decreased. While the changes in size varied across cell types, we observed that there was a more pronounced surround under natural movies compared to white noise for all cell types.

**Fig. 3.**
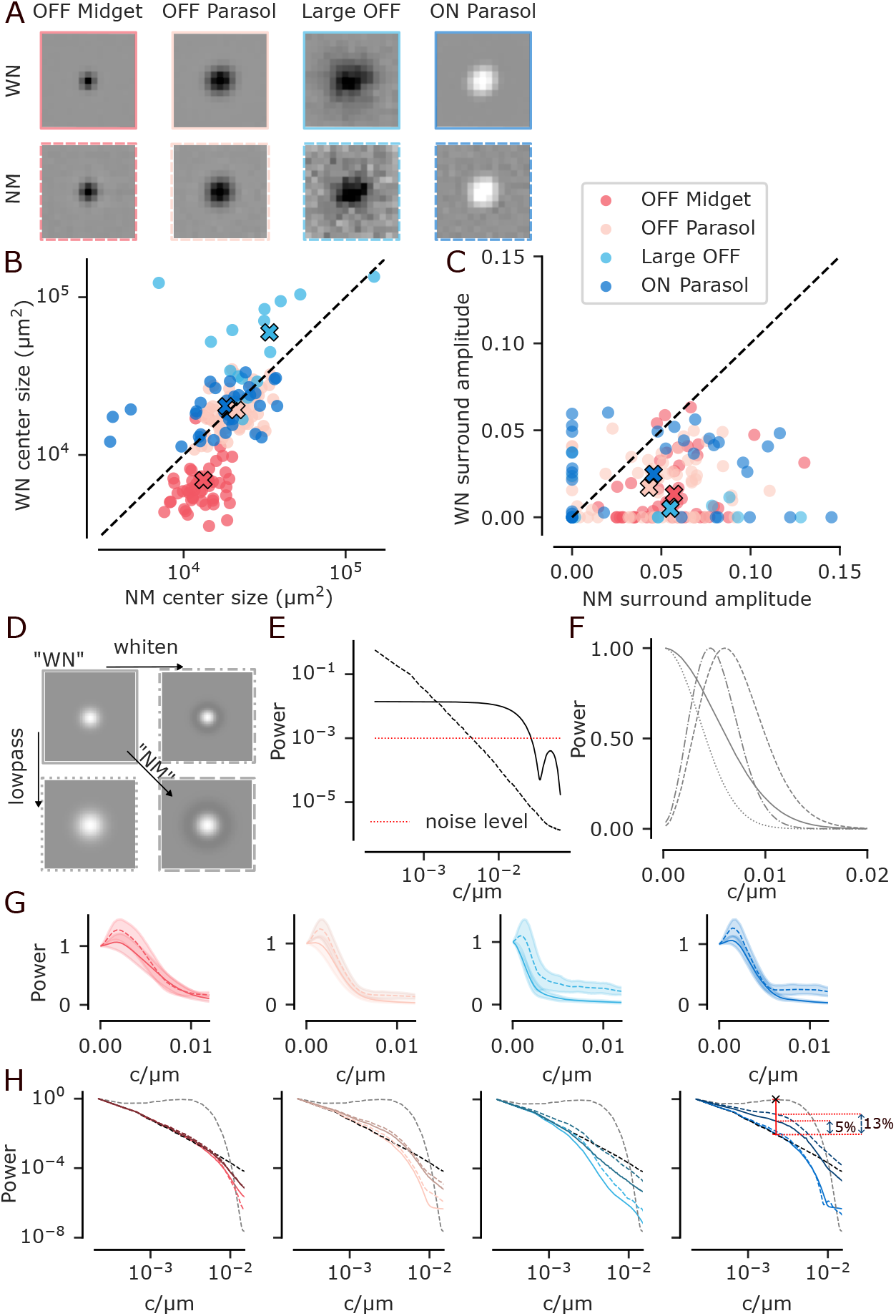
**A**. Cell-type-specific spatial RF comparison: Average spatial RF across neurons. First row: white noise. Second row: natural movies. **B**. Size of RF center for white noise vs. natural movies. **C**. Strength of RF surround strength for white noise vs. natural movies. Values below 1*e*−4 clipped to 0. **D**. Illustration of how size of RF center and magnitude of surround affect filter properties. Horizontal: surround contributes to whitening. Vertical: increased size contributes to low-pass filtering. The combination (bottom right) would be the prediction of efficient coding. **E**. Power spectrum of natural movie (black, dashed) and white noise stimulus (black, solid). Red dotted line: Flat spectrum of a hypothetical noise level. **F**. Transfer functions of the four theoretical filters from D. Line styles correspond to the outlines of the boxes in D. **G**. Average transfer functions of the cell-type-specific RFs from LN models trained on white noise (solid line) and natural movies (dashed) for OFF midget (first plot, red), OFF parasol (second plot, pink), large OFF (third plot, turquoise) and ON parasol cells (fourth plot, blue). Shaded regions: SD. **H**. Power spectrum ofthe original natural movie (black, dashed) with natural movie spectra after filtering with different filters: the simulated whitening filter (gray, dashed; panel E, bottom right), the rank-one model RFs estimated on movies (light coloured dashed lines) and on white noise (light coloured solid lines) and the rank-two model RFs estimated on movies (dark coloured dashed lines) and white noise (dark coloured solid lines) for different cell types. The red vertical line represents the distance between the original natural movie spectrum and a flat spectrum at the RF relevant frequency. White-noise, rank-two estimated RF flattens it by 5% (distance between first two dotted red lines), natural movie rank-two estimated RF flattens it by 13% (distance between first and third dotted red line). All spectra are normalized to the first data point.

To corroborate these results, we quantified the size and amplitude strength by fitting a difference of two elliptical Gaussians (DoG) to the spatial kernels. We calculated the size of the center as the area of the ellipse encompassing two standard deviations of the inner Gaussian fit. The amplitude of the surround was the absolute value of the minimum (maximum) of the DoG fit for an ON (OFF) cell, respectively. Our results (Figure 3B,C) show a significant increase in size for midget cells (Wilcoxon signed-rank test, *p* < .001 N= 51) and a significant increase in surround amplitude strength for all cell types (individual tests for cell types, Wilcoxon signed-rank test, OFF midget: *p* < .001 N=51; OFF parasols: *p* < .001 N=69; large OFF *p* = .005 N=16; ON parasol: *p* = .02 N=36).

### Surrounds contribute little to spatial whitening of the responses to natural stimuli

We asked what could explain these RF changes. They resembled changes predicted by efficient coding (Figure 3A). Based on the efficient coding hypothesis (Atick and Redlich, 1992), we can expect two ways in which the receptive fields should adjust between white noise and natural movie stimulation if the information about the stimulus being transmitted from the eye to the brain was to be maximized, considering constrained resources.

First, the filter for natural movies should have a center-surround structure to remove correlations which are present in natural environments. The noise stimulus is already white in the frequencies influenced by RGCs RFs and therefore the RFs should not show a surround (Figure 3D, horizontal).

Second, the RFs should be larger for natural movie stimuli compared to white noise stimuli because of the difference in power in high frequencies and the resulting signal-to-noise ratio (SNR) of the two stimuli (Figure 3D, vertical). Natural movies exhibit a 1/*f* ^2^ spectrum, where the power decreases steeply with frequency. Therefore, the SNR of the movie stimulus also drops faster with increasing frequencies, compared to the almost flat spectrum of the white noise stimulus. To illustrate this, we consider photon noise with a flat spectrum. When a stimulus power spectrum intersects with the photon noise spectrum, the SNR becomes too low for meaningful information extraction. For the natural movie spectrum, this intersection occurs at a lower frequency compared to the flat spectrum of the white noise stimulus (Figure 3E). As a result, the high frequency information becomes too unreliable in natural movies, resulting in larger filters with lower cutoff frequencies being optimal (Figure 3F, dashed). Conversely, white noise retains a sufficiently high SNR at higher frequencies, allowing for smaller filters with higher cutoff frequencies, therefore including higher-frequency information (Figure 3F, solid).

Midget cells were the only cell type where the predicted increase in the size of the center occurred comparing natural movie stimulation to white noise stimulation. The increase in surround strength under natural movie stimulation occurred for all cell types. We previously quantified the increase in the strength of the surround (Figure 3C). However, the question remains how much this increase in surround actually contributes to the whitening effect predicted by efficient coding. To quantify this, we filtered the natural movie signal using the LN-model-estimated RF filters. Then, we computed the spatial power spectra of the filtered signal and averaged them over the cells of each cell type. For all the cell types, the spectra filtered by RFs estimated from natural movies were not whitened stronger than those estimated from white noise. Overall, all of the RFs caused little to no whitening of the spectral content of natural movie stimulus.

### Rank-two model reveals some degree of whitening for ON parasol cells

A potential confound of our analysis could be that we assumed space-time separability for the receptive fields. Prior work suggests that space-time separated LN models may underestimate the surround’s strength due to its temporal delay with respect to the center (Kilavik, Silveira, and Kremers, 2003; Cowan, Sabharwal, and Wu, 2016). This delay cannot be captured by a single spatial and temporal filter pair. To control for this confound, we trained rank-two LN models which contain two independent spatial and temporal filter pairs for the center and surround. These models had similar predictive performance as the rank-one models on white noise with an average Pearson’s correlation of 0.69 (0.73) rank-two (rank-one) for OFF midgets; 0.7 (0.7) OFF parasols; 0.34 (0.34) large OFF cells; 0.64 (0.64) ON parasols. On natural movies the performances were also very similar between the two model types, namely 0.77 (0.76) OFF midgets; 0.77 (0.77) OFF parasols; 0.53 (0.52) large OFF cells; 0.87 (0.86) ON parasols. The rank-two models’ in-domain vs. out-of-domain performance also followed the previously observed results: best performance was achieved in-domain and they generalized better from white noise to natural stimuli then the other way around.

We repeated the filtering analysis with the RFs estimated by rank-two models. The whitening effect was negligible for all cell types except of ON parasol cells. For ON parasol cells, filters estimated from white noise indeed flattened the spectrum somewhat (by 5%) relative to a flat spectrum, while those estimated from natural movies achieved a stronger (13%) flattening (Figure 3H). In contrast, a flattening of 88% was achieved by synthetic filters which we optimized to whiten the movie signal while using the same constraints on receptive field size as the RGCs (Figure 3H). These numbers are reported for the spatial frequency at which the whitening effect was maximal within the relevant range given the RF sizes. Thus, ON parasol cells did whiten the spectra to some extent under both stimuli, more so under natural movies, qualitatively following the efficient coding hypothesis at least to some extent. On the other hand, three out four cell types we studied did not show an increase in whitening despite them exhibiting a stronger surround under natural movie stimulation. This result points towards the surround serving a different purpose than the one of efficient coding.

## Discussion

### LN models

In this paper we showed that LN models give rise to different response functions depending on which stimulus ensemble – white noise or natural scenes – they are trained on. A number of previous studies have argued for using natural scenes to study visual processing, because models trained on white noise stimuli do not generalize well to natural movie inputs (McIntosh et al., 2017; Sinz et al., 2018; Heitman et al., 2016). Our results differ from these earlier findings and extend them in several ways:

First, also in our hands, LN models trained on noise perform worse on movies compared to models directly trained on movies. However, unlike reported by Heitman et al. (2016), we find this performance gap to be fairly small. Second, we also observed the converse effect: Training models only on movies does not allow to accurately predict how RGCs respond to noise. This finding is in contrast to work in primary visual cortex, where the generalization from movies to noise was better than from noise to movies (Talebi and Baker, 2012; Sinz et al., 2018). However, we note that care is needed when interpreting such numbers. Overall, responses to movies were better predicted in our data than responses to noise, presumably because movies contain stronger local luminance fluctuations, which are “easy” to predict and account for a larger fraction of the total variance compared to noise stimuli. Thus, meaningfully interpreting absolute differences in predictive performance may be difficult. Generally, performance metrics based on explained variance are somewhat difficult to interpret. While only a perfect model would reach 100% performance, differences across models do not necessarily tell us how interesting the explained variance is in terms of the underlying computational phenomena it captured.

### Spatial Adaptation

Our findings demonstrate that RGCs adapt their spatial RFs in response to differences in stimulus spectra, specifically between spatiotemporal white noise and natural movie stimuli. We observed two types of changes – in the RF center sizes and in surround amplitude strength.

The observed RF size changes varied across cell types. OFF midget cells showed increased RF center sizes under natural movie stimulation. This increase is consistent with the expected adaptation to the lower SNR at high frequencies in natural movies compared to white noise, as can be derived from the efficient coding theory (Atick and Redlich, 1992). Parasol cells exhibited no significant size changes, potentially because their RFs are already large enough to effectively function as low-pass filters. Large OFF cells showed decreased RF sizes under natural movies, though these remained larger than parasol RFs. The increase in surround amplitude strength occurred for all cell types.

Similar changes have been reported in higher visual areas. Lesica et al. (2007) also found that in the cat LGN, the cells’ RFs increased both the center size and surround amplitude strength under natural stimuli compared to white noise stimulation. Given the resemblance of the adaptation we observed in the RGCs – the OFF midget cells specifically – it could originate in the retina. Sharpee, Sugihara, et al. (2006) demonstrated that RFs of cat V1 neurons adapt to stimulus statistics by tuning toward lower spatial frequencies under natural movie stimulation compared to white noise. This tuning aligns with the SNR adaptation predictions and matches the midget cells’ RF changes in our study.

### The efficient coding theory

There are two key aspects to consider regarding efficient coding theory in light of the RF structure and changes we show: (1) its general prediction about RF structure, such as the center-surround organization, and (2) its prediction about RF adaptation to specific stimulus statistics, such as stronger surrounds under natural movie stimulation.

The observed center-surround structure in ON parasol cells could, despite its low strength, support the idea that these cells have evolved to process the general statistics of the visual environment, in line with efficient coding. Their RFs may be sub-optimal for white noise but are well-suited to partly decorrelate natural scenes by whitening the signal (Atick and Redlich, 1992). However, the extent of whitening in the ON parasol cells – 13% (5%) when filtering with movie (noise) estimated filters relative to a fully whitened i.e. flat spectrum – and the absence of any significant whitening in the other three cell types suggests that the efficient coding principle may not fully explain their RF structure.

The RF surround is not the only mechanism by which the retina could achieve whitening of natural stimuli. Multiple studies have suggested other decorrelation mechanisms, such as a nonlinear transformation at the output stage of the retina (Pitkow and Meister, 2012; Turner, Schwartz, and Rieke, 2018) and fixational eye movements (Kuang et al., 2012; Segal et al., 2015). Therefore, it is still possible that the retina whitens the input signal for efficient encoding. For this reason, we do not invalidate the efficient coding theory as a whole, our contribution lies in establishing that the presence of a surround in the RF does not automatically translate to whitening.

A linear receptive field that whitens also does not necessarily directly translate to decorrelated cell responses. Important to consider is the non-linearity of the RF itself. While classical LN models consider the center and surround as additive components, recent studies have shown that the interplay between both components could also be multiplicative or involve more complex nonlinearities (Turner and Rieke, 2016) and these non-linearities can evoke correlated activity (Karamanlis, Khani, et al., 2025).

Regarding RF adaptation across stimulus statistics, the midget cells did increase in size as would be predicted by efficient coding. And while the other cell types did not, it could be, as stated above, due to their RF already be large enough to be effective low-pass filters. With respect to the surround, only ON parasol cells demonstrated a stronger surround that could contribute to whitening under natural movies compared to white noise, consistent with the efficient coding theory predicting stimulus-specific encoding. Yet, the white noise filter for ON parasol cells produced a stronger whitening effect than the natural movie filters of all other cell types. This observation suggests that it is not necessarily the change in stimulus that is responsible for the center-surround RF structure. Rather, it appears that ON parasol cells have a more whitening receptive field compared to the other cell types, independent of stimulus statistics.

### Temporal adaptation

Nirenberg et al. (2010) focused on temporal adaptation between white noise and natural movie stimuli of RGCs in the retina. We did not study temporal adaptation for two reasons. Firstly, we pool multiple retinas, and the temporal profiles of cells of the same cell types are not well comparable across retinas (Zhao et al., 2019). Secondly, the natural movie stimulus was presented at a 25% lower mean luminance than the white noise stimulus, which could additionally affect the temporal properties of the cells (Benardete and Kaplan, 1999).

While the changes in luminance also affect the spatial properties of RGCs RFs, it cannot account for the adaptation we observed. Lower luminance tends to increases the size of the RF, but this change generally occurs at a threshold when switching from predominantly cone to predominantly rod vision (Troy, Bohnsack, and Diller, 1999). Neither of our stimuli is dark enough to trigger this switch. Lower luminance also decreases the strength of the surround (Thoreson and Mangel, 2012; Farrow et al., 2013). In our case, however, the stimulus with the higher surround strength is the one with a lower luminance. Therefore, it is safe to assume that the surround that appears does not do so because of a luminance decrease but due to the differences in stimulus statistics.

## Acknowledgment

This project was funded by the Deutsche Forschungsgemeinschaft (DFG, German Research Foundation) - Project-ID 432680300 - SFB 1456 and the European Research Council (ERC) under the European Union’s Horizon Europe research and innovation programme (Grant agreement No. 101041669). The authors acknowledge the computing time made available on the high-performance computers HLRN-IV at GWDG at the NHR Center NHR@Göttingen. The center is jointly supported by the Federal Ministry of Education and Research and the state governments participating in the NHR (www.nhr-verein.de/unsere-partner). M.F.B thanks the International Max Planck Research School for Intelligent Systems (IMPRS-IS).

## Data availability

All data and code will be made available upon publication.

